# Regional and age-dependent changes in ubiquitination in cellular and mouse models of Spinocerebellar ataxia type 3

**DOI:** 10.1101/2023.02.01.526671

**Authors:** Haiyang Luo, Sokol V. Todi, Henry L. Paulson, Maria do Carmo Costa

## Abstract

Spinocerebellar ataxia type 3 (SCA3), also known as Machado–Joseph disease, is the most common dominantly inherited ataxia. SCA3 is caused by a CAG repeat expansion in the *ATXN3* gene that encodes an expanded tract of polyglutamine (polyQ) in the disease protein ataxin-3 (ATXN3). As a deubiquitinating enzyme, ATXN3 regulates numerous cellular processes including proteasome- and autophagy-mediated protein degradation. In SCA3 disease brain, polyQ-expanded ATXN3 accumulates with other cellular constituents, including ubiquitin (Ub)-modified proteins, in select areas like the cerebellum and the brainstem, but whether pathogenic ATXN3 affects the abundance of ubiquitinated species is unknown. Here, in mouse and cellular models of SCA3, we investigated whether elimination of murine *Atxn3* or expression of wild-type or polyQ-expanded human ATXN3 alters soluble levels of overall ubiquitination, as well as K48-linked (K48-Ub) and K63-linked (K63-Ub) chains. Levels of ubiquitination were assessed in the cerebellum and brainstem of 7- and 47-week-old *Atxn3* knockout and SCA3 transgenic mice, and also in relevant mouse and human cell lines. In older mice, we observed that wild-type ATXN3 impacts the cerebellar levels of K48-Ub proteins. In contrast, pathogenic ATXN3 leads to decreased brainstem abundance of K48-Ub species in younger mice and changes in both cerebellar and brainstem K63-Ub levels in an age-dependent manner: younger SCA3 mice have higher levels of K63-Ub while older mice have lower levels of K63-Ub compared to controls. Human SCA3 neuronal progenitor cells also show a relative increase in K63-Ub proteins upon autophagy inhibition. We conclude that wild-type and mutant ATXN3 differentially impact K48-Ub- and K63-Ub-modified proteins in the brain in a region- and age-dependent manner.

## 2 Introduction

Spinocerebellar ataxia type 3 (SCA3), also known as Machado–Joseph disease, is the most common form of dominantly inherited ataxia (Schols et al., 2004; Paulson, 2007; Costa Mdo and Paulson, 2012). Patients with SCA3 display progressive ataxia as a core symptom accompanied by other neurological signs (Coutinho and Andrade, 1978; Paulson, 2007) that reflect neuronal dysfunction and loss in the cerebellum, brainstem, and spinal cord (Seidel et al., 2012b). SCA3 is caused by expansion of a polyglutamine (polyQ)-encoding CAG trinucleotide repeat in the *ATXN3* gene (Kawaguchi et al., 1994), which encodes the ataxin-3 protein (ATXN3). The *ATXN3* (CAG)_n_ repeat length ranges from 12 to 44 in healthy individuals and from ~60 to 87 in SCA3 carriers (Maciel et al., 2001; Lima et al., 2005).

ATXN3 is a ubiquitously expressed deubiquitinating enzyme (DUB) that regulates many cellular processes including proteasome and autophagy-mediated protein degradation, transcription, cytoskeletal dynamics, and DNA damage repair (Costa Mdo and Paulson, 2012; Da Silva et al., 2019; Matos et al., 2019). While polyQ-expanded ATXN3 accumulates with other proteins, including ubiquitin (Ub)-modified proteins in selective areas of the brain (Chai et al., 1999; Chai et al., 2001; Chai et al., 2004; Seidel et al., 2012b), whether its ability to handle ubiquitinated proteins differs from wild-type ATXN3 is not known.

Posttranslational modification of proteins with Ub enables complex cellular signaling codes. Through isopeptide bonds, for example, the amino groups of lysine (K) or of initial methionine residues can be monoubiquitinated, multi-monoubiquitinated, homotypically-polyubiquitinated (e.g., K48- or K63-linked Ub chains), or heterotypically-polyubiquitinated (mixed chains, branched chains, combination of Ub and Ub-like proteins, with additional spatial complications added by the modification of Ub itself by phosphorylation, etc.) (Dikic and Schulman, 2022). In addition, hydroxyl groups in serine and threonine residues (via ester bonds), and thiol groups in cysteine (C) residues (via thioester bonds) can be ubiquitinated (Pao et al., 2018; Kelsall et al., 2019; Mabbitt et al., 2020). Further increased complexity to the Ub code stems from Ub linkage modification by sugars and lipids (Yang et al., 2017; Chatrin et al., 2020; Otten et al., 2021; Kelsall et al., 2022).

This complex Ub “language”, comprising different Ub modifications and chains that encode distinct cellular signals, involves code “writers” (E3 ligases in combination with E1 and E2 enzymes that add Ub to proteins), “readers” (molecules harboring Ub-binding domains that distinguish specific Ub modifications and target the ubiquitinated substrate to a downstream process) and “erasers” (DUBs that reverse ubiquitination) (Dikic and Schulman, 2022). DUBs recognizing specific Ub modifications therefore provide an important layer of regulation of the Ub code and cellular function.

ATXN3 displays DUB isopeptidase, esterase, thioesterase (Burnett et al., 2003; De Cesare et al., 2021), and deneddylase activities (Ferro et al., 2007). Given its broad molecular network, ATXN3 may also possess other types of protease activities (Costa Mdo and Paulson, 2012; Matos et al., 2019). ATXN3 contains a globular, amino-terminal, Josephin domain harboring the catalytic residues and two Ub-binding sites, and a carboxyl-terminal domain of largely undetermined structure containing two Ub interacting motifs (UIMs) followed by the polyQ tract and a third UIM, depending on the isoform (Goto et al., 1997; Masino et al., 2003; Nicastro et al., 2005; Nicastro et al., 2010). The catalytic cysteine residue (C14) in the Josephin domain is essential for protease activity, and the UIMs are required for selective polyUb binding (Burnett et al., 2003; Chai et al., 2004; Mao et al., 2005; Nicastro et al., 2005; Winborn et al., 2008).

Although ATXN3 favors hydrolysis of isopeptide bonds from polyUb chains with four or more Ub molecules *in vitro* (Burnett et al., 2003; Winborn et al., 2008), it also deubiquitinates specific monoubiquitinated substrates (Scaglione et al., 2011). Among polyUb chains, ATXN3 favors *in vitro* deubiquitination of K63-linked and K48/K63-mixed chains over K48-linked chains (Winborn et al., 2008). K48-linked chains (K48-Ub) primarily target proteins for proteasomal degradation (Dikic and Schulman, 2022) and K63-linked polyUb (K63-Ub) signals for the formation of inclusion bodies and initiation of autophagy, among other roles (Tan et al., 2008; Chen et al., 2019; Blount et al., 2020). While overall levels of soluble protein ubiquitination are increased in brains of *Atxn3* knockout mice (Scaglione et al., 2011), whether this increase represents a preferential *in vivo* activity of ATXN3 towards a specific type of ubiquitination, and whether the expanded polyQ tract affects ATXN3 activity, are unknown.

Here, we investigate whether mouse *Atxn3* ablation and wild-type or polyQ-expanded human ATXN3 impact soluble levels of overall ubiquitination and K48-Ub and K63-Ub polyubiquitinated chains in mouse and cellular models of disease.

## 3 Materials and Methods

### 3.1 Animals

All animal procedures were approved by the University of Michigan Institutional Animal Care and Use Committee and conducted in accordance with the U.S. Public Health Service’s Policy on Humane Care and Use of Laboratory Animals (protocols PRO00008397 and PRO00008409). *Atxn3* knockout (Atxn3 KO) (Reina et al., 2010), and SCA3 YACMJD15.4 and YACMJD84.2 transgenic mice (Cemal et al., 2002) (all in a C57Bl/6J background) were genotyped using tail biopsy DNA isolated prior to weaning and genotypes were confirmed postmortem, as previously described (Reina et al., 2010; do Carmo Costa et al., 2013). For biochemical analysis, animals were euthanized with a lethal dose of ketamine-xylazine and PBS-perfused and brain regions were macro-dissected for biochemical assessments, as previously described (Reina et al., 2010; do Carmo Costa et al., 2013; Ashraf et al., 2019).

### 3.2 Cell culture and treatment

Mouse embryonic fibroblasts (MEFs) from both Atxn3 KO mice and WT littermates (Reina et al., 2010) were maintained in DMEM with 10% FBS, 1% Non-essential Amino Acids solution and 1% penicillin/streptomycin at 37□ and 5% CO2. Control and SCA3 neuronal progenitor cells (NPCs) were generated from the respective human embryonic stem cell (hESC) lines (CTRL, NIH registry #0147; SCA3, NIH registry # 0286), and were maintained in STEMdiff Neural Progenitor Medium (NPM), as previously described (Ashraf et al., 2020). Where indicated, cells were treated with lactacystin (15μM; Enzo Life Sciences), Chloroquine (100μM; Sigma-Aldrich), or DMSO (Sigma-Aldrich) for 12 hours.

### 3.3 Immunoblotting

Total proteins were extracted in radioimmunoprecipitation assay (RIPA) buffer from macro-dissected cerebellar and brainstem tissues, as previously described (do Carmo Costa et al., 2013) from Atxn3 KO (Reina et al., 2010; Toulis et al., 2020), SCA3 transgenic YACMJD15.4 and YACMJD84.2 (Cemal et al., 2002) and their wild-type mouse littermates (N=3 males or females per genotype), and stored at −80°C. Mouse brain protein lysates were resolved in 4–20% gradient sodium dodecyl sulfate-polyacrylamide (SDS)-PAGE electrophoresis gels and transferred to 0.45 μm PVDF membranes. Membranes were incubated overnight at 4°C with various antibodies: rabbit anti-pan ubiquitin (1:1,000, #3933S; Cell Signaling Technology, Danvers, MA), rabbit anti-K48-linkage specific polyUb (D9D5) (1:1,000, #8081; Cell Signaling Technology, Danvers, MA), rabbit anti-K63-linkage specific polyUb (D7A11) (1:1,000, #5621S; Cell Signaling Technology, Danvers, MA), mouse anti-ATXN3 (1H9) (1:1,000, MAB5360; Millipore, Billerica, MA), and mouse anti-GAPDH (1:50,000, MAB374; Millipore, Billerica, MA). Bound primary antibodies were visualized by incubation with peroxidase-conjugated anti-mouse or anti-rabbit secondary antibodies (1:10,000; Jackson Immuno Research Laboratories, West Grove, PA) followed by treatment with ECL-plus reagent (Western Lighting; PerkinElmer, Waltham, MA) and exposure to autoradiography films. Band intensities were quantified using ImageJ analysis software (NIH, Bethesda, MD).

### 3.4 Statistical analyses

Comparison of groups was performed using Student’s t-test or One-way ANOVA with a post hoc Tukey test. All statistical analyses were performed using Prism 9. Statistical significance was considered at *p* value ≤ 0.05.

## 4 Results

### 4.1 Wild-type mouse Atxn3 impacts the abundance of high molecular weight K48-linked species in select mouse brain areas

We first evaluated whether wild-type murine Atxn3 impacts the brain levels of all ubiquitinated proteins and of proteins modified with K48-Ub or K63-Ub chains. By immunoblotting, we assessed the abundance of proteins that are ubiquitinated (pan-Ub) or modified specifically with K48-Ub or K63-Ub linkages in soluble protein lysates from cerebellum and brainstem (Seidel et al., 2012b) of *Atxn3* knockout (Atxn3 KO) mice (Reina et al., 2010) and their control littermates (WT). The cerebellum and brainstem were chosen because they are the two most heavily affected brain regions in SCA3.

Because disease manifestations of SCA3 typically appear in the third or fourth decade of life (Costa Mdo and Paulson, 2012) and ATXN3 DUB activity may differ with age, we evaluated the above Ub species in young (7-week-old) and aged (47-week-old) mice. For each immunoblot readout of pan-Ub, K48-Ub or K63-Ub-modified proteins, we quantified three groups: total proteins (entire lane), HMW proteins (≥250 KDa) and LMW proteins (<250 KDa).

While cerebellar levels of pan-Ub (Figure 1A) and K63-Ub (Figure 1C) were similar between Atxn3 KO and WT controls in young and old mice, the abundance of cerebellar HMW K48-Ub species in 47-week-old Atxn3KO mice showed a 70% increase compared to controls of the same age (Figure 1B).

**Figure 1.**
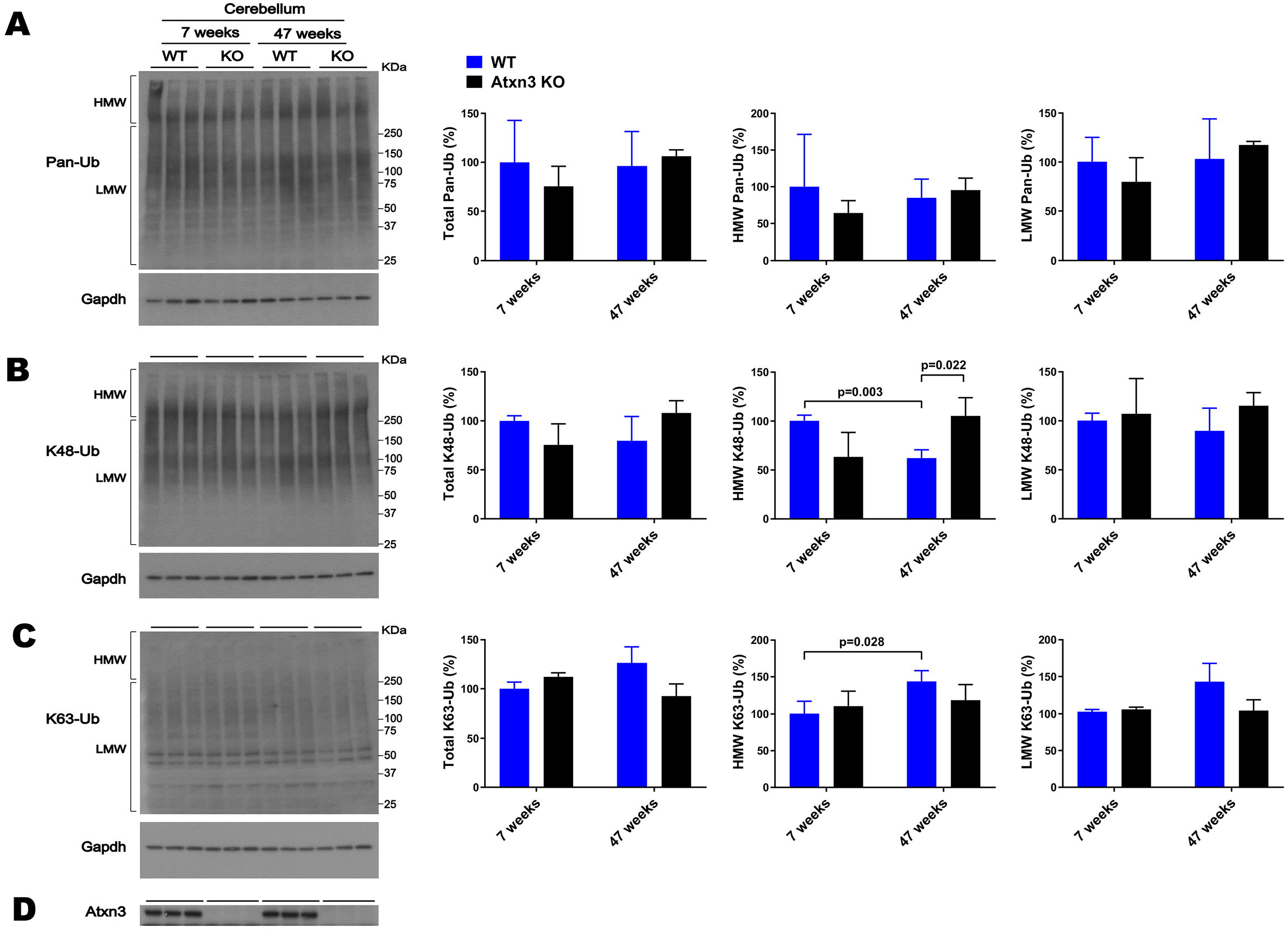
Cerebella of 47-week-old Atxn3 KO mice show increased levels of HMW K48-Ub species. Western blots and quantifications of Pan-Ub **(A)**, K48-Ub **(B)**, and K63-Ub **(C)** signal from the cerebella of 7- and 47-week-old Atxn3 KO mice and WT littermates. Graph bars represent the mean percentage (± SEM) of each Ub type normalized for Gapdh, relative to the levels of 7-week-old WT mice. *p* values are from Student’s t-test. **(D)** Immunoblot detecting mouse Atxn3 in the same mouse protein extracts.

Brainstem abundance of pan-Ub, K48-Ub, and K63-Ub was similar between Atxn3 KO and WT controls for both ages (Figure 2A-C), except for total pan-Ub amount, which was modestly decreased in 7-week-old Atxn3 KO compared to control mice (Figure 2A).

**Figure 2.**
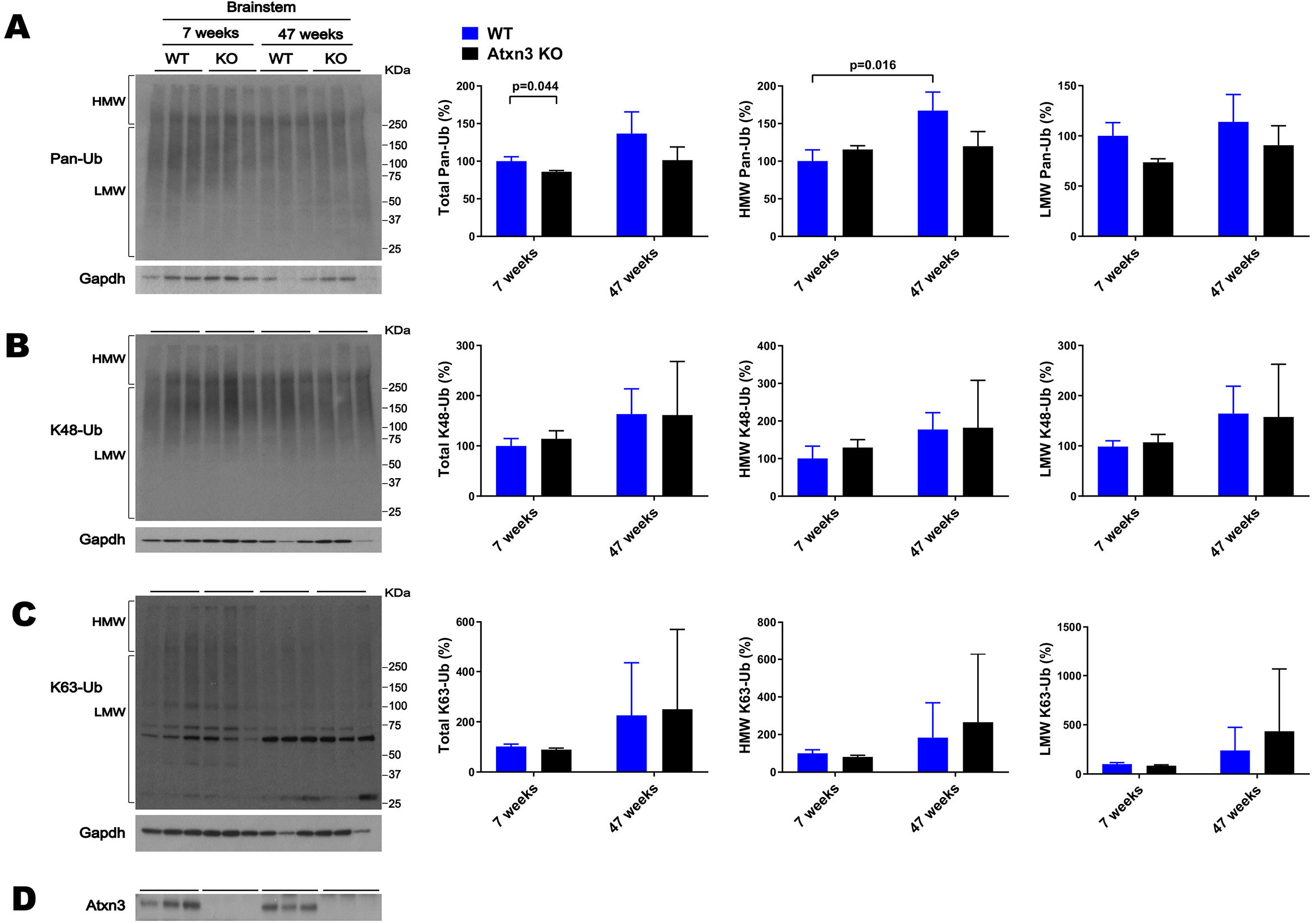
Brainstem of 7 and 47-week-old Atxn3 KO and WT littermate mice show similar levels of Pan-Ub, K48-Ub and K63-Ub species. Western blots and quantifications of Pan-Ub **(A)**, K48-Ub **(B)**, and K63-Ub **(C)** signal from the brainstems of 7- and 47-week-old Atxn3 KO mice and WT littermates. Graph bars represent the mean percentage (± SEM) of each Ub type normalized for Gapdh relative to the levels of 7- week-old WT mice. *p* values are from Student’s t-test. **(D)** Immunoblot detecting mouse Atxn3 in the same mouse protein extracts.

In summary, absence of *Atxn3* in the brain led to differences in some Ub species, but not others, implying a potentially selective regional and age-dependent activity of this DUB in overall Ub physiology.

We also observed differences in HMW polyUb species as a function of age: a) cerebella of 47-week-old WT mice displayed a 40% decrease in HMW K48-Ub (Figure 1B) and a 45% increase in HMW K63-Ub (Figure 1C) compared to 7-week-old mice; and b) brainstems of 47-week-old WT mice showed about 70% more HMW Pan-Ub than 7-week-old mice (Figure 2A). As expected (Dulka et al., 2021), aging is associated with regional differences in the types of Ub species in brains of wild-type mice.

### 4.2 Atxn3 affects levels of HMW K48-Ub proteins in mouse embryonic fibroblasts upon proteasomal inhibition

The above results suggest that Atxn3 regulates levels of HMW K48-Ub species in the aged mouse brain, potentially in a region-specific manner (Figure 1B). K48-Ub chains mediate proteasomal degradation (Dikic and Schulman, 2022) and proteasomal activity is known to decrease with aging (Schmidt and Finley, 2014). Accordingly, we assessed whether levels of K48-Ub are impacted by loss of Atxn3 in non-brain cells subjected to proteasomal inhibition, which impacts levels of K48-Ub (Dikic and Schulman, 2022).

Similar to our results in aged Atxn3 KO mouse cerebella, MEFs derived from Atxn3 KO mice (Reina et al., 2010) treated with the proteasome inhibitor lactacystin selectively showed an approximately 20% increase in HMW K48-Ub species compared to WT controls (Figure 3B). Mirroring our results in mouse brain, an absence of Atxn3 in MEFs did not affect levels of pan-Ub or K63-Ub species, whether in vehicle- (DMSO) or lactacystin-treated MEFs (Figure 3A,C). However, Atxn3 KO MEFs treated with DMSO did show a 35% decrease of total K48-Ub compared with MEFs derived from WT littermates (Figure 3B), an effect that was not observed in baseline conditions (data not shown). This latter observation suggests that Atxn3-mediated effects on K48-Ub changes are impacted by DMSO, which is known to alter cellular conditions in a concentration-dependent manner (Braden and Neufeld, 2016; Dludla et al., 2018; Bhatt et al., 2020). Moreover, proteasomal inhibition in Atxn3 KO and WT MEFs led to similar increases in the levels of all types of K63-Ub proteins (Figure 3C) confirming that K63-Ub species also signal for proteasomal degradation (Ohtake et al., 2018).

**Figure 3.**
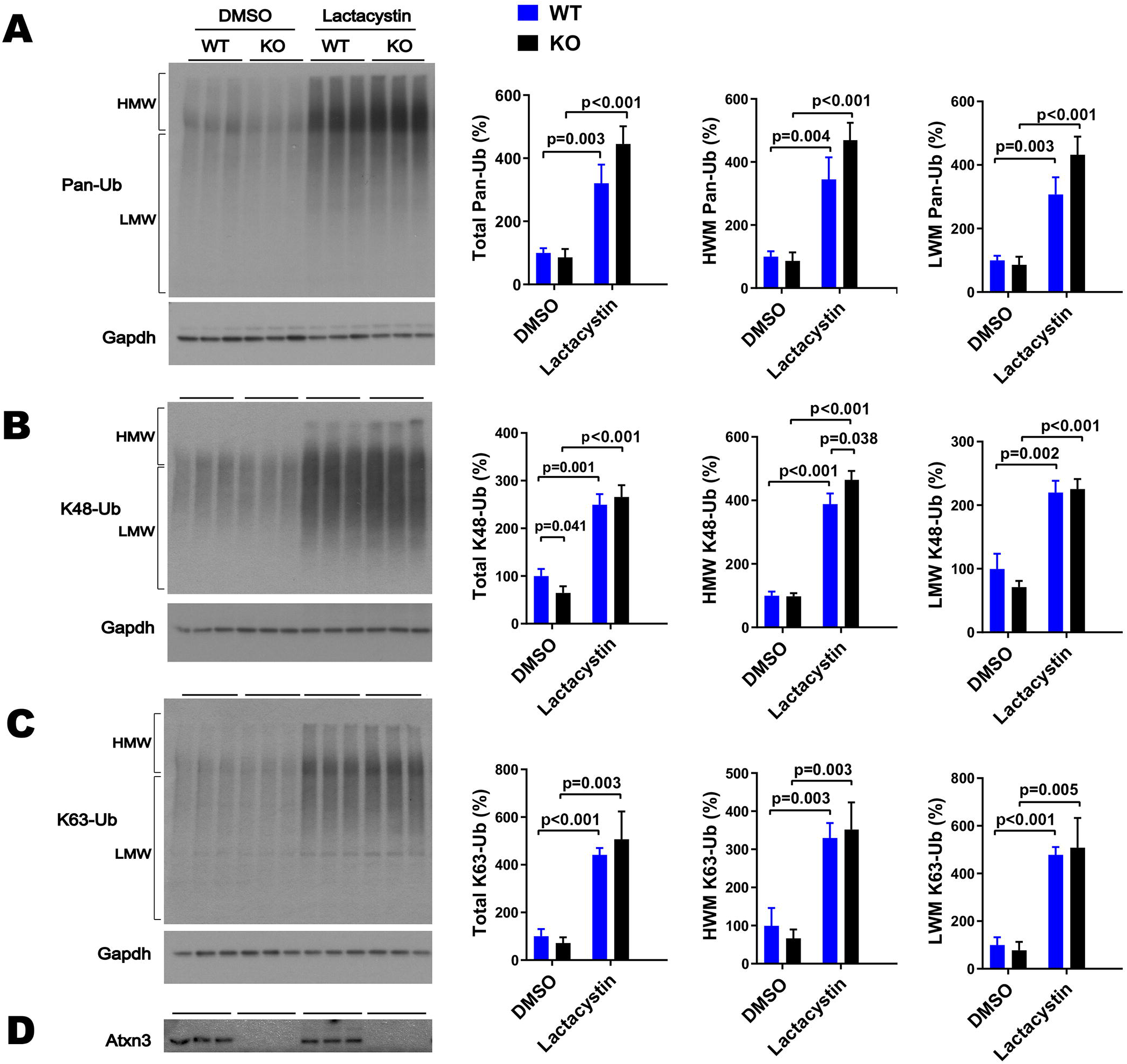
Levels of pan-Ub, K48-Ub and K63-Ub species in Atxn3 KO and WT MEFs upon proteasomal inhibition. Westerns blot and quantifications of Pan-Ub **(A)**, K48-Ub **(B)**, and K63-Ub **(C)** signal from Atxn3 KO and WT MEFs treated with the proteasome inhibitor lactacystin or with vehicle (DMSO). Graph bars represent the mean percentage (± SEM) of each Ub type normalized for Gapdh relative to the levels of WT MEFs treated with DMSO. *p* values are from Student’s t-test. **(D)** Immunoblot detecting mouse Atxn3 in the same cell protein extracts.

### 4.3 Expression of polyQ-expanded human ATXN3 increases overall levels of ubiquitination in select brain areas of young SCA3 mice

We next evaluated whether exogenous overexpression of wild-type or polyQ-expanded human ATXN3 affects overall protein ubiquitination in the cerebellum and brainstem. We again assessed 7-week-old and 47-week-old mice, comparing mice overexpressing normal or pathogenic ATXN3 to WT littermates. SCA3 YACMJD transgenic mice express the full-length human *ATXN3* gene harboring a CAG repeat in either the nonpathogenic range (YACMJD15.4 mice) or the disease range (YACMJD84.2 mice) (Cemal et al., 2002). While both homozygous YACMJD84.2 (Q84/Q84) and hemizygous YACMJD84.2 (Q84) mice demonstrate intranuclear accumulation of ATXN3 in brain by 8 weeks of age, they differ phenotypically: Q84/Q84 mice display robust motor signs as early as 6 weeks of age whereas Q84 mice only show motor defects at ~75 weeks of age (do Carmo Costa et al., 2013). In contrast, hemizygous YACMJD15.4 (Q15) do not show motor impairment at any tested age, despite robustly overexpressing nonpathogenic ATXN3 (Cemal et al., 2002).

Whereas 7-week-old WT, Q15 and Q84 mice exhibited similar cerebellar levels of pan-Ub (Figure 4A), 7-week-old Q84/Q84 mice showed a 60-110% increase in cerebellar abundance of total, HMW and LMW pan-Ub compared to WT mice (Figure 4A). In contrast, 47-week-old mice showed no detectable differences in cerebellar pan-Ub levels across all genotypes (WT, Q15, Q84, and Q84/Q84; Figure 4A).

**Figure 4.**
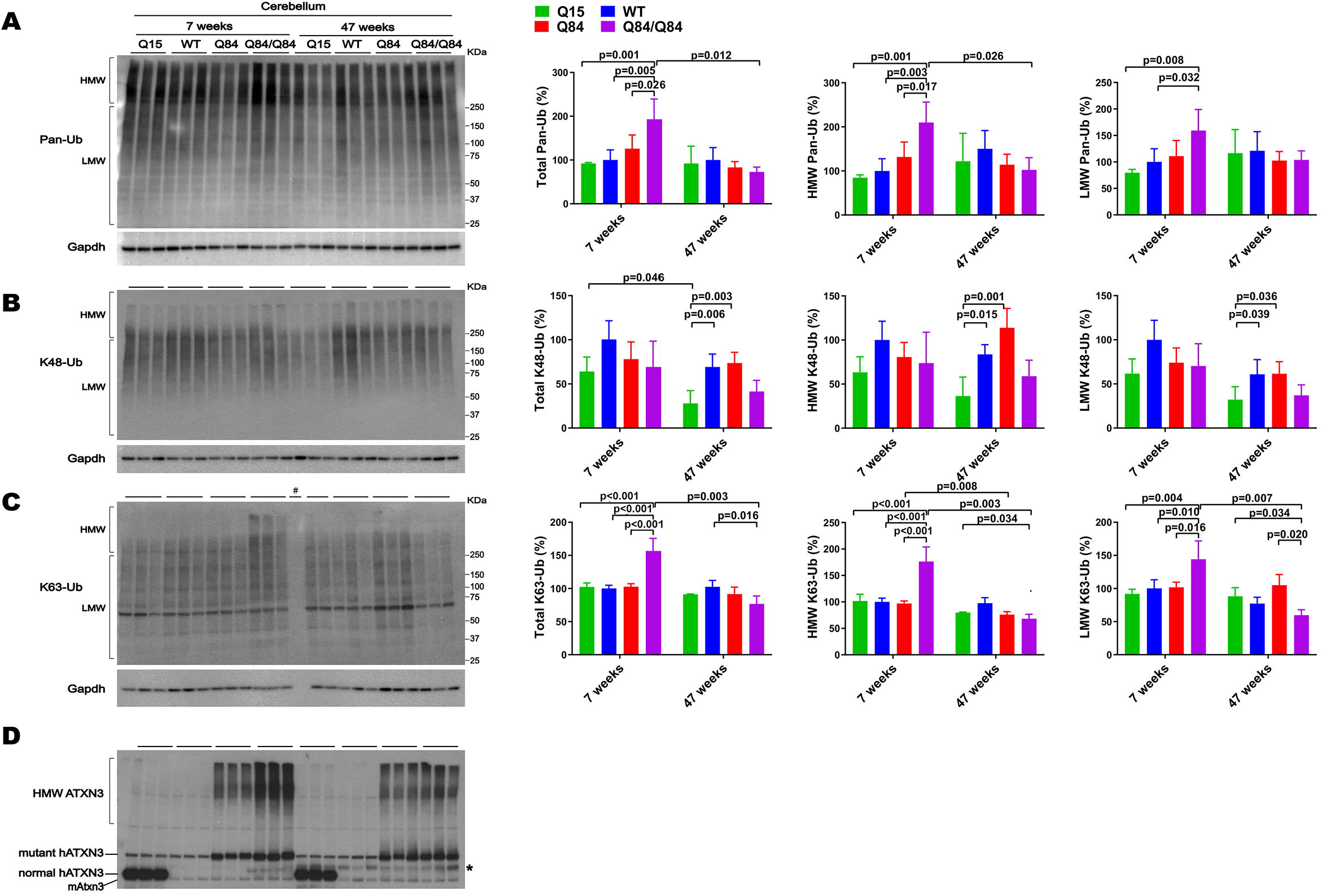
Cerebella of young homozygous SCA3 transgenic (Q84/Q84) mice show increased Pan-Ub and K63-Ub species. Westerns blot quantification of Pan-Ub **(A)**, K48-Ub **(B)**, and K63-Ub **(C)** modified proteins in cerebella of 7- and 47-week-old hemizygous Q15, hemizygous Q84, homozygous Q84/Q84, and WT littermate mice. Graph bars represent the mean percentage (± SEM) of each Ub type normalized for Gapdh relative to the levels of 7-week-old WT mice. *p* values are from one-way ANOVA with a post hoc Tukey test. **(D)** Immunoblot detecting mouse and human ATXN3 in the same mouse protein extracts. *Represents band corresponding to nonspecific detection of mouse IgG.

Brainstem levels of pan-Ub were similar in all four genotypes at 7 and 47 weeks of age (Figure 5A) except that HMW pan-Ub levels in 7-week-old Q84 mice were 39% lower than in Q84/Q84 mice (Figure 5A). Furthermore, in older mice across all genotypes, brainstem levels of pan-Ub were lower (Figure 5A).

**Figure 5.**
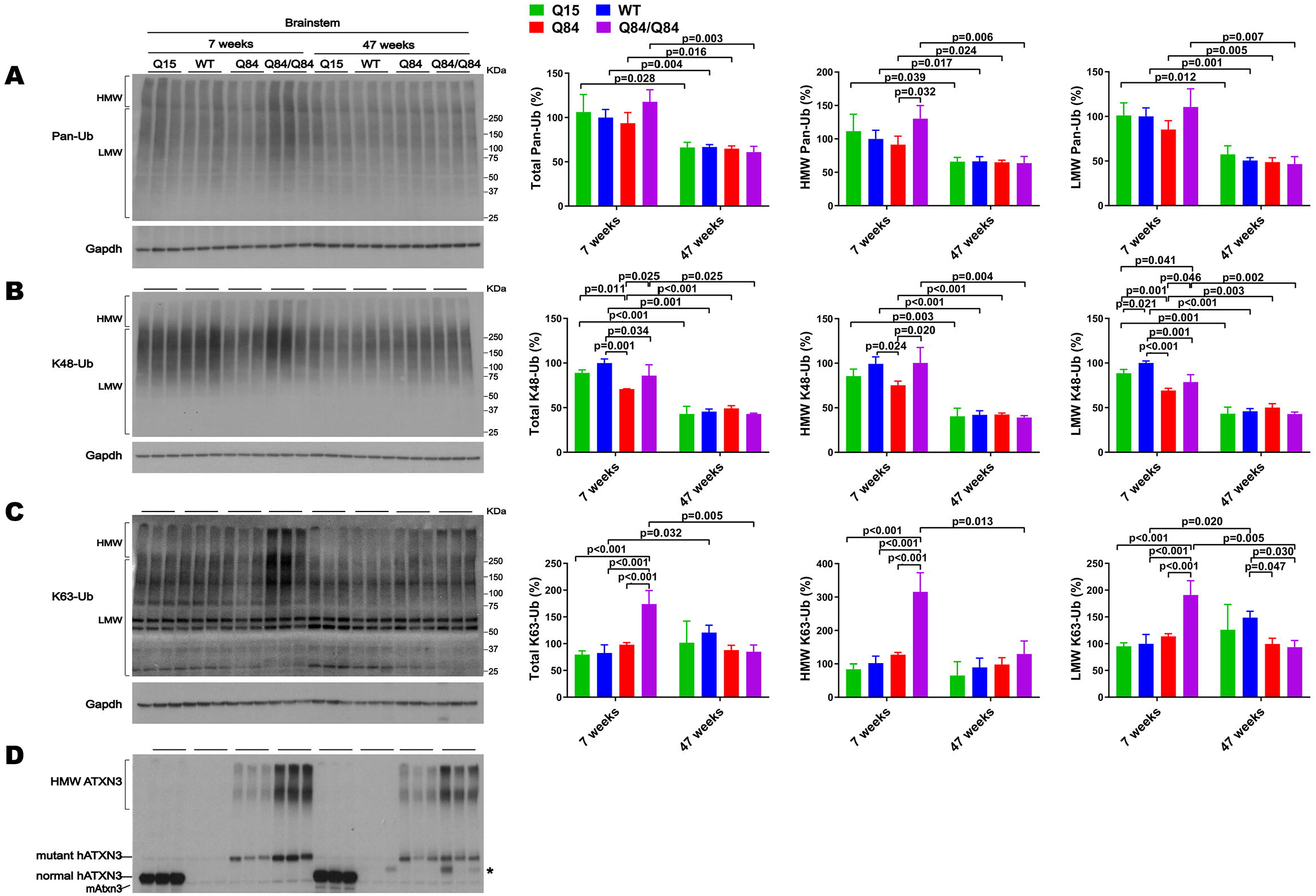
Brainstem of young homozygous Q84/Q84 mice show marked increase in K63-Ub species and a slight decrease in K48-Ub species. Western blots and quantifications of Pan-Ub **(A)**, K48-Ub **(B)**, and K63-Ub **(C)** signal from the brainstems of 7- and 47-week-old hemizygous Q15, hemizygous Q84, homozygous Q84/Q84, and WT littermate mice. Graph bars represent the mean percentage (± SEM) of each Ub type normalized for Gapdh relative to the levels of 7-week-old WT mice. *p* values are from One-way ANOVA with a pos hoc Tukey test. **(D)** Immunoblot detecting mouse and human ATXN3 in the same mouse protein extracts. *Represents nonspecific detection of mouse IgG.

### 4.4 Wild-type and polyQ-expanded human ATXN3 differentially impact brain K48-linked and K63-linked polyubiquitination in SCA3 mice

We subsequently evaluated whether K48-linked ubiquitination is affected by expression of wild-type or polyQ-expanded human ATXN3 in young versus aged SCA3 transgenic mice. In 7-week-old mice, cerebellar abundance of K48-Ub linkages was similar across all four genotypes (Figure 4B). In 47-week-old mice, however, K48-Ub species were decreased by approximately 40% in the cerebellum, but not brainstem, of Q15 mice compared to WT and hemizygous Q84 mice (Figure 4B). This result suggests that, with aging, overexpression of wild-type human ATXN3 modulates the abundance of K48-Ub chains in the cerebellum. At 7 weeks of age, brainstem K48-Ub levels were decreased by 20-30% in Q84 (total, MHW, and LMW) and Q84/Q84 (total and LMW) mice compared to WT mice (Figure 5B). At 47 weeks of age, however, no differences in brainstem K48-Ub levels were observed across the various genotypes (Figure 5B). Yet as with pan-Ub, at 47 weeks of age for all genotypes, levels of K48-Ub were decreased compared to that in mice at 7 weeks of age (Figure 5B).

K63-Ub abundance patterns revealed a robust, age-dependent change in homozygous Q84/Q84 mice (Figure 4C and Figure 5C). Cerebella and brainstems of 7-week-old Q84/Q84 mice displayed a striking 50-200% increase in the levels of total, HMW and LMW K63-Ub compared with other groups (Figure 4C and Figure 5C). Remarkably, and in contrast to results in younger mice, 47-week-old Q84/Q84 mice showed a 20-37% decrease in LMW K63-Ub levels in the cerebellum and brainstem, compared to WT mice (Figure 4C and Figure 5C). Brainstems of 47-week-old hemizygous Q84 mice also showed a 35% decrease in LMW K63-Ub compared to controls (Figure 5C). These results suggest that pathogenic ATXN3 triggers a spike in the overall abundance of K63-Ub species in early stages of motor dysfunction, followed by a modest decrease of some K63-Ub species in later stages of disease.

### 4.5 Human SCA3 neuronal progenitor cells display an increase in overall ubiquitinated species at baseline and increased accumulation of K63-Ub signal upon autophagy inhibition

The SCA3 mouse lines used above overexpress wild-type or polyQ-expanded human ATXN3. To probe whether physiological levels of expanded ATXN3 affect the abundance of Ub species in human neuronal cells, we assessed the levels of Ub species in neuronal progenitor cells (NPCs) derived from a hESC line harboring one disease and one healthy allele (SCA3, NIH registry # 0286) (Moore et al., 2019; Ashraf et al., 2020). For comparison, we also assessed NPCs derived from a hESC line harboring two nonpathogenic *ATXN3* alleles (CTRL, NIH registry #0147).

Under basal conditions, SCA3 NPCs displayed a 75% increase in HMW pan-Ub levels compared to CTRL NPCs (Figure 6A). This difference was maintained upon proteasomal inhibition, but not upon autophagy inhibition (Figure 6A).

**Figure 6.**
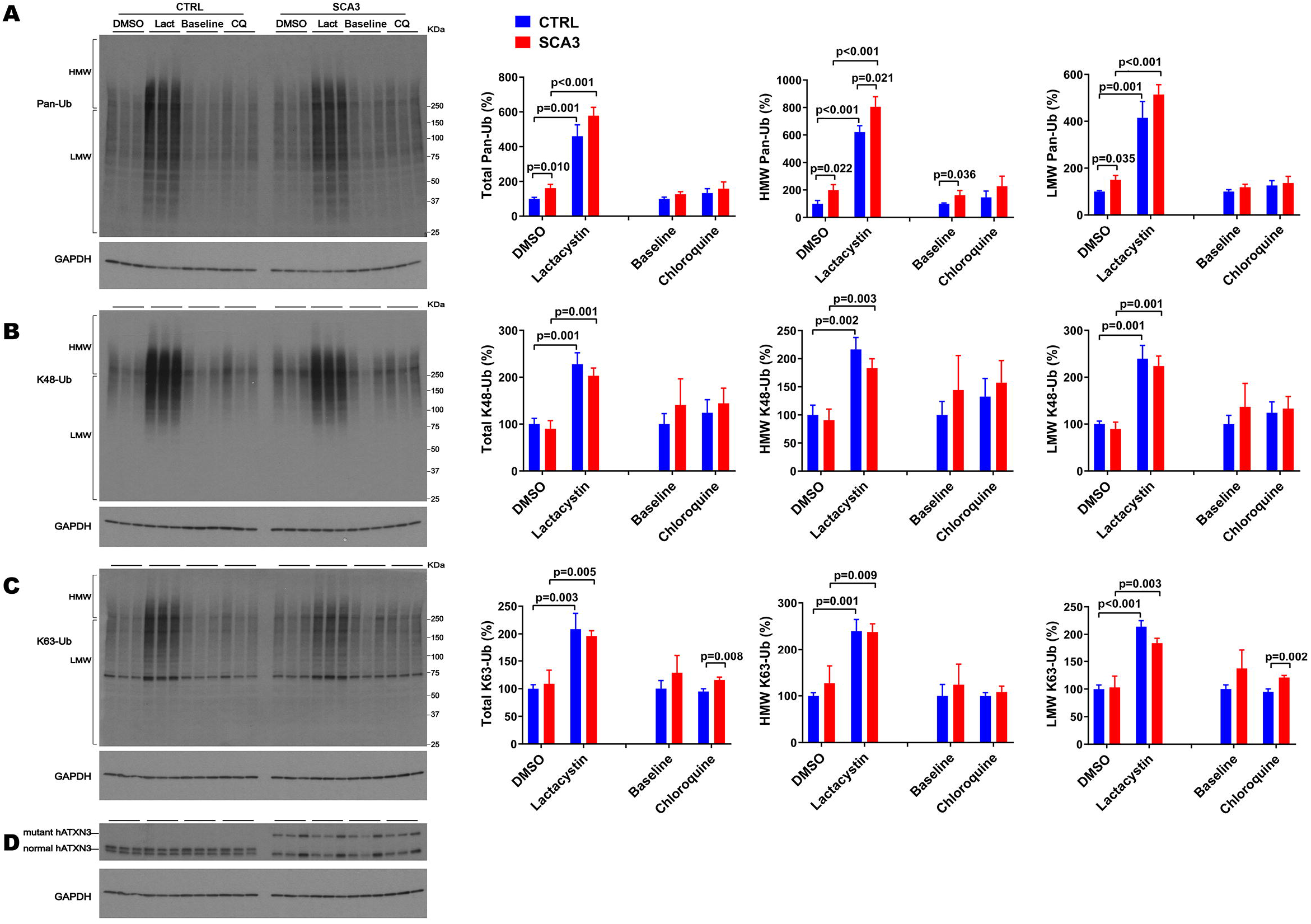
SCA3 NPCs display increased overall ubiquitinated species in basal conditions and increased K63-Ub proteins upon autophagy inhibition. Western blots and quantifications of Pan-Ub **(A)**, K48-Ub **(B)**, and K63-Ub **(C)** signal from proteins in SCA3 and CTRL neuronal progenitor cells (NPCs) in basal conditions (baseline), treated with lactacystin (Lact, proteasome inhibitor) or vehicle (DMSO), or treated with chloroquine (CQ, autophagy inhibitor). Graph bars represent the mean percentage (± SEM) of each Ub type normalized for Gapdh relative to the levels of CTRL NPCs treated with DMSO. *p* values are from Student’s t-test. **(D)** Immunoblot detecting human ATXN3 in the same cell protein extracts.

With respect to linkage-specific Ub species, total and LMW K63-Ub were increased by 25% in SCA3 NPCs compared to CTRL NPCs upon autophagy inhibition (Figure 6C). In contrast, no differences in K48-Ub were noted between SCA3 and CTRL NPCs under any conditions (Figure 6B).

## 5 Discussion

Ub-positive intracellular aggregates have been recognized as a common feature of polyQ diseases for more than two decades (Jana and Nukina, 2003), but how Ub metabolism is affected in polyQ disorders is poorly understood. Assessment of the landscape of specific Ub protein modifications has been quite limited in polyQ diseases other than Huntington’s disease (HD) (Bennett et al., 2007). The accumulation of Ub, proteasomal subunits, and p62 in ATXN3-positive aggregates in human SCA3 disease brain and cell and mouse models (Paulson et al., 1997; Schmidt et al., 1998; Chai et al., 1999; Cemal et al., 2002; Seidel et al., 2012a) implies that Ub metabolism is dysregulated in SCA3. Since ATXN3 is a DUB, such impairment could reflect altered DUB function of pathogenic ATXN3 due to conformational changes in the protein, or to its sequestration into aggregates. Because the Ub code is complex and ATXN3 displays versatile DUB activity, several forms of Ub protein modifications could be affected in SCA3. Here, we explored whether wild-type and mutant ATXN3 affect the abundance of pan-Ub, K48-Ub, and K63-Ub chains. Figure 7 summarizes the differences observed in Ub species in mouse models. The abundance of pan-Ub, K48-Ub and K63-Ub in two vulnerable brain regions, the cerebellum and brainstem, is differentially affected by eliminating mouse *Atxn3* or by overexpressing wild-type or mutant human ATXN3, with specific changes in Ub species occurring in a region- and age-dependent manner. Two key findings are: 1) wild-type ATXN3 itself influences the abundance of HMW K48-Ub species in a region- and age-dependent manner; and 2) pathogenic, polyQ-expanded ATXN3 affects the levels of linkage-specific Ub species in an age- and region-specific manner. In addition, as previously reported in the mouse cortex (Bennett et al., 2007), we observed that the abundance of total pan-Ub and K48-Ub, but not K63-Ub, decreases with age in the cerebellum and brainstem of all the SCA3 transgenic mouse groups we examined (Figure 7), confirming that levels of certain types of Ub protein modifications change with aging.

**Figure 7.**
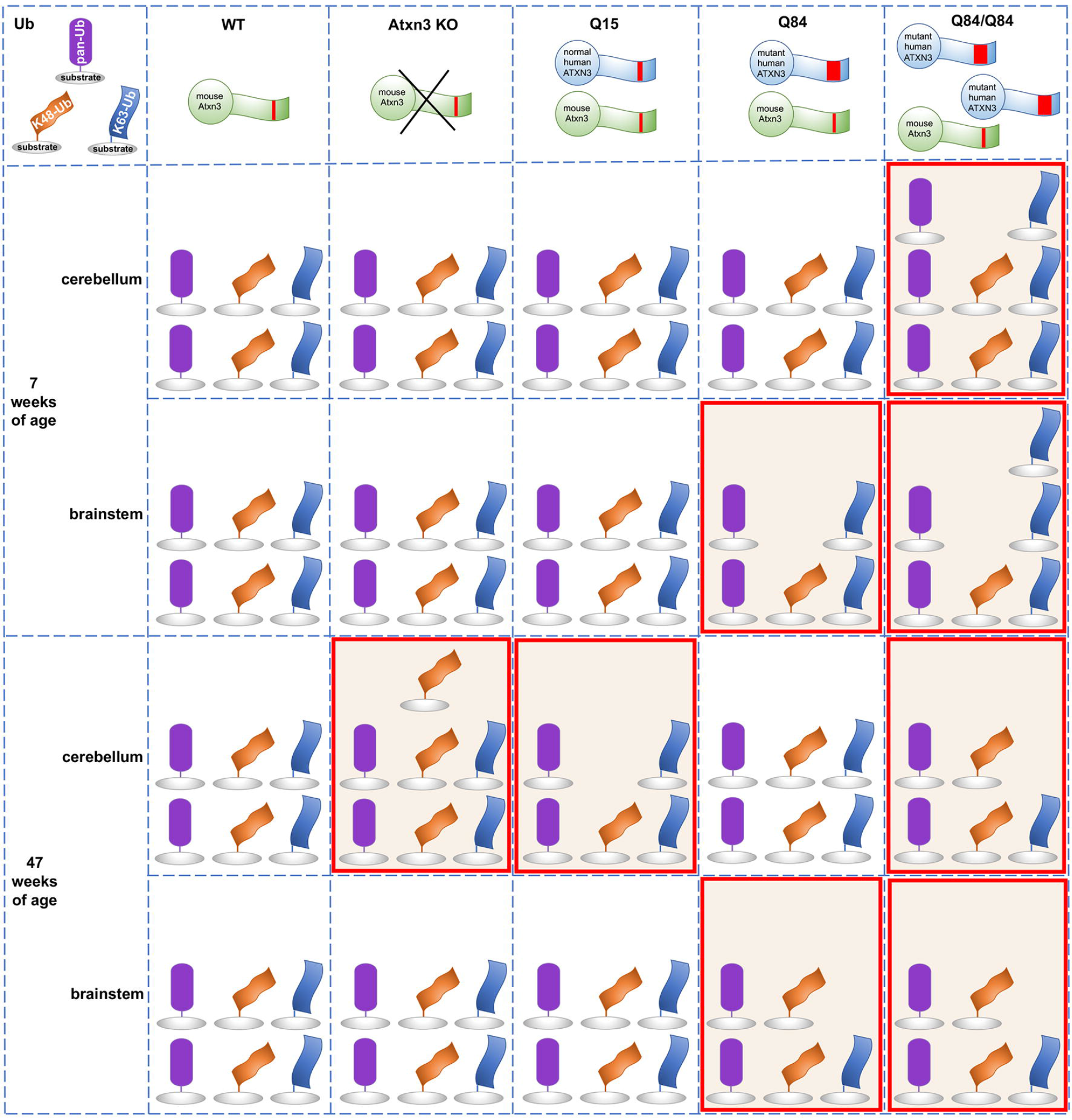
Summary of the abundance of pan-Ub, K48-Ub and K63-Ub species in the cerebellum and the brainstem of 7 and 47-week-old Atxn3 KO and SCA3 transgenic mice.

An earlier study of Atxn3 KO mice reported higher levels of HMW pan-Ub species in whole brain lysates (Scaglione et al., 2011). We instead observed similar pan-Ub levels in the cerebellum and brainstem of young and aged Atxn3 KO mice compared to wild-type controls (Figures 7). This apparent discrepancy between the two studies may reflect region-specific differences in Ub signaling. Consistent with our results in mice, MEFs lacking Atxn3 show increased levels of HMW K48-Ub species upon proteasomal inhibition (Figure 3B), in line with the reported *in vitro* preference of ATXN3 for cleaving longer Ub chains (Burnett et al., 2003). K48-Ub chains target proteins for proteasomal degradation (Dikic and Schulman, 2022) and aging is associated with reduced or impaired proteasomal function (Chondrogianni et al., 2003). Our results suggest that wild-type ATXN3, a stress-response protein (Reina et al., 2012), participates in the regulation of K48-Ub species that accumulate when the proteasome is impaired (Bennett et al., 2007), such us in the cerebella of aged (47-week-old) mice. While recombinant wild-type ATXN3 shows modest preference *in vitro* for short K63-Ub linked chains over short K48-Ub linked polyUb (Winborn et al., 2008), we noted no change in the abundance of soluble K63-Ub proteins in the cerebellum and brainstem of Atxn3 KO mice versus wild type or Q15 mice, or in MEFs lacking or containing Atxn3 (Figure 7). Thus, any preference that wild-type ATXN3 shows for K63-Ub linkages likely depends on the complex physiological context of the cell, which is not recapitulated by reconstituted Ub chain cleavage studies *in vitro*.

Expression of polyQ-expanded ATXN3 robustly increases the abundance of soluble K63-Ub proteins in the cerebellum and brainstem of SCA3 mice at a young age (7-weeks) when they already display motor dysfunction (Figure 7). Similarly, the cortex and striatum of HD R6/2 transgenic mice expressing a pathogenic huntingtin fragment display increased levels of K63-Ub chains (Bennett et al., 2007). While K63-linked Ub changes may prove to be a common theme among polyQ diseases, further work assessing Ub chain composition in other polyQ disorders is clearly needed.

K63-linked ubiquitination is implicated in specific cell signaling pathways. Because one role of K63-Ub chains is to signal proteins for autophagy (Blount et al., 2020) and pathogenic ATXN3 is reported to impair autophagy initiation (Ashkenazi et al., 2017), the accumulation of soluble K63-Ub in the cerebellum and brainstem of young Q84/Q84 mice may indicate that polyQ-expanded ATXN3 impedes autophagy initiation at early stages of disease. Further evidence supporting the hypothesis that mutant ATXN3 promotes an accumulation of soluble K63-Ub proteins comes from our observation of increased levels of K63-Ub in SCA3 NPCs, but not CTRL NPCs, upon autophagy inhibition (Figure 6C), and from an earlier report demonstrating impaired autophagy and the accumulation of large autophagosome-like vesicles in SCA3 disease brain (Sittler et al., 2017). Given this possibility, why do we observe *decreased* soluble K63-Ub in the cerebellum and brainstem of aged Q84/Q84 mice (Figure 7)? Conceivably, K63-Ub proteins become sequestered in insoluble ATXN3 aggregates in later stages of disease and therefore are relatively depleted in the soluble protein lysates that we analyzed in this study.

While polyQ-expanded ATXN3 appears to markedly affect the abundance of soluble K63-Ub species in the two assessed vulnerable brain areas in SCA3, it selectively increases levels of soluble global Ub in the cerebellum, but not brainstem, of homozygous Q84/Q84 mice at early stages of overt motor impairment (Figure 7) and in SCA3 NPCs under basal and proteasomal inhibition conditions (Figure 6A). In contrast, young and aged hemizygous Q84 and WT littermates show similar abundance of soluble Ub proteins in the cerebellar and brainstem (Figure 7), paralleling the reported findings of unchanged global Ub in the cortex and striatum of HD patients and mice (Bennett et al., 2007). These results suggest that pathogenic ATXN3 selectively impacts global ubiquitination levels in cells undergoing cellular stress.

With respect to K48-Ub, pathogenic ATXN3 was associated with modestly and selectively decreased K48-Ub levels in the brainstem of 7-week-old hemizygous Q84 and homozygous Q84/Q84 mice (Figure 7). Because wild-type ATXN3 impacts K48-Ub abundance when the proteosome is compromised (Figures 1–3 and 7), our data in SCA3 mice suggest that pathogenic ATXN3 may normally modulate soluble K48-Ub levels in brain regions of SCA3 mice potentially undergoing proteasomal stress due to its own expression. However, the impact of polyQ-expanded ATXN3 on soluble K48-Ub levels appears to be modest and transient, only noticeable in early stages of SCA3 disease progression. In contrast, overexpression of a pathogenic huntingtin fragment sustainably increased the amount of K48-Ub linkages throughout disease progression (Bennett et al., 2007). Overall, polyQ-expanded ATXN3 and the pathogenic huntingtin fragment appear to show common and distinct effects in the abundance of specific Ub chain modified proteins, suggesting that the protein context together with the polyQ tract drives their specific impact in Ub metabolism. Future work is required to explicate similarities and differences in the Ub landscape among polyQ disorders.

Limitations of the current study include the fact that we captured the ubiquitin environment at only two time points, using antibodies that provide a low-resolution view of the ubiquitin landscape in the central nervous system. Moreover, we focused our attention on two brain regions vulnerable in SCA3, the cerebellum and brainstem, without assessing other brain regions, including basal ganglia, cerebral cortex and spinal cord, that show variable involvement. More comprehensive studies are needed to determine whether the region- and age-specific alterations in the Ub landscape reported here also occur in other brain regions over time. In addition, extending the analysis beyond soluble Ub species to insoluble material may shed light on the relationship between disease protein aggregation and changes to the Ub environment.

In summary, we report Ub linkage-, regional- and age-dependent differences in mouse brain and relevant mammalian cells caused by the expression of wild type or pathogenic ATXN3. These findings provide further physiological context for the normal functions of ATXN3 in the mammalian brain and will aid future investigations of substrates and cellular pathways regulated by this disease-linked DUB.

## 7 Author contributions

HL, SVT, HLP and MCC contributed to the conception and the design of the study. HL conducted the experiments, organized and analyzed the data, and prepared the figures. MCC provided the mouse brain materials and cell lines, supervised the experimental work, analyzed the data, and wrote the manuscript. SVT, HLP and MCC wrote and edited the manuscript.

## 9 Acknowledgements

The authors would like to thank Ilya Bezprozvanny for providing the YACMJD84.2 mice.

## 10 Data availability statement

The original contributions presented in the study are included in the article/supplementary material, further inquiries can be directed to the corresponding authors.

## References

Ashkenazi, A., Bento, C.F., Ricketts, T., Vicinanza, M., Siddiqi, F., Pavel, M., et al. (2017). Polyglutamine tracts regulate beclin 1-dependent autophagy. Nature 545(7652), 108–111. doi: 10.1038/nature22078.

Ashraf, N.S., Duarte-Silva, S., Shaw, E.D., Maciel, P., Paulson, H.L., Teixeira-Castro, A., et al. (2019). Citalopram Reduces Aggregation of ATXN3 in a YAC Transgenic Mouse Model of Machado-Joseph Disease. Mol Neurobiol 56(5), 3690–3701. doi: 10.1007/s12035-018-1331-2.

Ashraf, N.S., Sutton, J.R., Yang, Y., Ranxhi, B., Libohova, K., Shaw, E.D., et al. (2020). Druggable genome screen identifies new regulators of the abundance and toxicity of ATXN3, the Spinocerebellar Ataxia type 3 disease protein. Neurobiol Dis 137, 104697. doi: 10.1016/j.nbd.2019.104697.

Bennett, E.J., Shaler, T.A., Woodman, B., Ryu, K.Y., Zaitseva, T.S., Becker, C.H., et al. (2007). Global changes to the ubiquitin system in Huntington’s disease. Nature 448(7154), 704–708. doi: 10.1038/nature06022.

Bhatt, D., Stan, R.C., Pinhata, R., Machado, M., Maity, S., Cunningham-Rundles, C., et al. (2020). Chemical chaperones reverse early suppression of regulatory circuits during unfolded protein response in B cells from common variable immunodeficiency patients. Clin Exp Immunol 200(1), 73–86. doi: 10.1111/cei.13410.

Blount, J.R., Johnson, S.L., and Todi, S.V. (2020). Unanchored Ubiquitin Chains, Revisited. Front Cell Dev Biol 8, 582361. doi: 10.3389/fcell.2020.582361.

Braden, C.R., and Neufeld, T.P. (2016). Atg1-independent induction of autophagy by the Drosophila Ulk3 homolog, ADUK. FEBS J 283(21), 3889–3897. doi: 10.1111/febs.13906.

Burnett, B., Li, F., and Pittman, R.N. (2003). The polyglutamine neurodegenerative protein ataxin-3 binds polyubiquitylated proteins and has ubiquitin protease activity. Hum Mol Genet 12(23), 3195–3205. doi: 10.1093/hmg/ddg344 [pii].

Cemal, C.K., Carroll, C.J., Lawrence, L., Lowrie, M.B., Ruddle, P., Al-Mahdawi, S., et al. (2002). YAC transgenic mice carrying pathological alleles of the MJD1 locus exhibit a mild and slowly progressive cerebellar deficit. Hum Mol Genet 11(9), 1075–1094.

Chai, Y., Berke, S.S., Cohen, R.E., and Paulson, H.L. (2004). Poly-ubiquitin binding by the polyglutamine disease protein ataxin-3 links its normal function to protein surveillance pathways. J Biol Chem 279(5), 3605–3611. doi: 10.1074/jbc.M310939200 [pii],

Chai, Y., Koppenhafer, S.L., Shoesmith, S.J., Perez, M.K., and Paulson, H.L. (1999). Evidence for proteasome involvement in polyglutamine disease: localization to nuclear inclusions in SCA3/MJD and suppression of polyglutamine aggregation in vitro. Hum Mol Genet 8(4), 673–682. doi: ddc073 [pii].

Chai, Y., Wu, L., Griffin, J.D., and Paulson, H.L. (2001). The role of protein composition in specifying nuclear inclusion formation in polyglutamine disease. J Biol Chem 276(48), 44889–44897. doi: 10.1074/jbc.M106575200[pii].

Chatrin, C., Gabrielsen, M., Buetow, L., Nakasone, M.A., Ahmed, S.F., Sumpton, D., et al. (2020). Structural insights into ADP-ribosylation of ubiquitin by Deltex family E3 ubiquitin ligases. Sci Adv 6(38). doi: 10.1126/sciadv.abc0418.

Chen, R.H., Chen, Y.H., and Huang, T.Y. (2019). Ubiquitin-mediated regulation of autophagy. J Biomed Sci 26(1), 80. doi: 10.1186/s12929-019-0569-y.

Chondrogianni, N., Stratford, F.L., Trougakos, I.P., Friguet, B., Rivett, A.J., and Gonos, E.S. (2003). Central role of the proteasome in senescence and survival of human fibroblasts: induction of a senescence-like phenotype upon its inhibition and resistance to stress upon its activation. J Biol Chem 278(30), 28026–28037. doi: 10.1074/jbc.M301048200.

Costa Mdo, C., and Paulson, H.L. (2012). Toward understanding Machado-Joseph disease. Prog Neurobiol 97(2), 239–257. doi: S0301-0082(11)00212-7 [pii] 10.1016/j.pneurobio.2011.11.006.

Coutinho, P., and Andrade, C. (1978). Autosomal dominant system degeneration in Portuguese families of the Azores Islands. A new genetic disorder involving cerebellar, pyramidal, extrapyramidal and spinal cord motor functions. Neurology 28(7), 703–709.

Da Silva, J.D., Teixeira-Castro, A., and Maciel, P. (2019). From Pathogenesis to Novel Therapeutics for Spinocerebellar Ataxia Type 3: Evading Potholes on the Way to Translation. Neurotherapeutics 16(4), 1009–1031. doi: 10.1007/s13311-019-00798-1.

De Cesare, V., Carbajo Lopez, D., Mabbitt, P.D., Fletcher, A.J., Soetens, M., Antico, O., et al. (2021). Deubiquitinating enzyme amino acid profiling reveals a class of ubiquitin esterases. Proc Natl Acad Sci U S A 118(4). doi: 10.1073/pnas.2006947118.

Dikic, I., and Schulman, B.A. (2022). An expanded lexicon for the ubiquitin code. Nat Rev Mol Cell Biol, 1–15. doi: 10.1038/s41580-022-00543-1.

Dludla, P.V., Jack, B., Viraragavan, A., Pheiffer, C., Johnson, R., Louw, J., et al. (2018). A dose-dependent effect of dimethyl sulfoxide on lipid content, cell viability and oxidative stress in 3T3-L1 adipocytes. Toxicol Rep 5, 1014–1020. doi: 10.1016/j.toxrep.2018.10.002.

do Carmo Costa, M., Luna-Cancalon, K., Fischer, S., Ashraf, N.S., Ouyang, M., Dharia, R.M., et al. (2013). Toward RNAi Therapy for the Polyglutamine Disease Machado-Joseph Disease. Mol Ther 21(10), 1898–1908. doi: mt2013144 [pii] 10.1038/mt.2013.144.

Dulka, B.N., Trask, S., and Helmstetter, F.J. (2021). Age-Related Memory Impairment and Sex-Specific Alterations in Phosphorylation of the Rpt6 Proteasome Subunit and Polyubiquitination in the Basolateral Amygdala and Medial Prefrontal Cortex. Front Aging Neurosci 13, 656944. doi: 10.3389/fnagi.2021.656944.

Ferro, A., Carvalho, A.L., Teixeira-Castro, A., Almeida, C., Tome, R.J., Cortes, L., et al. (2007). NEDD8: a new ataxin-3 interactor. Biochim Biophys Acta 1773(11), 1619–1627. doi: S0167-4889(07)00191-7 [pii] 10.1016/j.bbamcr.2007.07.012.

Goto, J., Watanabe, M., Ichikawa, Y., Yee, S.B., Ihara, N., Endo, K., et al. (1997). Machado-Joseph disease gene products carrying different carboxyl termini. Neurosci Res 28(4), 373–377. doi: S0168-0102(97)00056-4 [pii],

Jana, N.R., and Nukina, N. (2003). Recent advances in understanding the pathogenesis of polyglutamine diseases: involvement of molecular chaperones and ubiquitin-proteasome pathway. J Chem Neuroanat 26(2), 95–101. doi: 10.1016/s0891-0618(03)00029-2.

Kawaguchi, Y., Okamoto, T., Taniwaki, M., Aizawa, M., Inoue, M., Katayama, S., et al. (1994). CAG expansions in a novel gene for Machado-Joseph disease at chromosome 14q32.1. Nat Genet 8(3), 221–228. doi: 10.1038/ng1194-221.

Kelsall, I.R., McCrory, E.H., Xu, Y., Scudamore, C.L., Nanda, S.K., Mancebo-Gamella, P., et al. (2022). HOIL-1 ubiquitin ligase activity targets unbranched glucosaccharides and is required to prevent polyglucosan accumulation. EMBO J 41(8), e109700. doi: 10.15252/embj.2021109700.

Kelsall, I.R., Zhang, J., Knebel, A., Arthur, J.S.C., and Cohen, P. (2019). The E3 ligase HOIL-1 catalyses ester bond formation between ubiquitin and components of the Myddosome in mammalian cells. Proc Natl Acad Sci U S A 116(27), 13293–13298. doi: 10.1073/pnas.1905873116.

Lima, M., Costa, M.C., Montiel, R., Ferro, A., Santos, C., Silva, C., et al. (2005). Population genetics of wild-type CAG repeats in the Machado-Joseph disease gene in Portugal. Hum Hered 60(3), 156–163.

Mabbitt, P.D., Loreto, A., Dery, M.A., Fletcher, A.J., Stanley, M., Pao, K.C., et al. (2020). Structural basis for RING-Cys-Relay E3 ligase activity and its role in axon integrity. Nat Chem Biol 16(11), 1227–1236. doi: 10.1038/s41589-020-0598-6.

Maciel, P., Costa, M.C., Ferro, A., Rousseau, M., Santos, C.S., Gaspar, C., et al. (2001). Improvement in the molecular diagnosis of Machado-Joseph disease. Arch Neurol 58(11), 1821–1827. doi: noc10138 [pii].

Mao, Y., Senic-Matuglia, F., Di Fiore, P.P., Polo, S., Hodsdon, M.E., and De Camilli, P. (2005). Deubiquitinating function of ataxin-3: insights from the solution structure of the Josephin domain. Proc Natl Acad Sci U S A 102(36), 12700–12705. doi: 0506344102 [pii] 10.1073/pnas.0506344102.

Masino, L., Musi, V., Menon, R.P., Fusi, P., Kelly, G., Frenkiel, T.A., et al. (2003). Domain architecture of the polyglutamine protein ataxin-3: a globular domain followed by a flexible tail. FEBS Lett 549(1-3), 21–25. doi: S0014579303007488 [pii],

Matos, C.A., de Almeida, L.P., and Nobrega, C. (2019). Machado-Joseph disease/spinocerebellar ataxia type 3: lessons from disease pathogenesis and clues into therapy. J Neurochem 148(1), 8–28. doi: 10.1111/jnc.14541.

Moore, L.R., Keller, L., Bushart, D.D., Delatorre, R.G., Li, D., McLoughlin, H.S., et al. (2019). Antisense oligonucleotide therapy rescues aggresome formation in a novel spinocerebellar ataxia type 3 human embryonic stem cell line. Stem Cell Res 39, 101504. doi: 10.1016/j.scr.2019.101504.

Nicastro, G., Menon, R.P., Masino, L., Knowles, P.P., McDonald, N.Q., and Pastore, A. (2005). The solution structure of the Josephin domain of ataxin-3: structural determinants for molecular recognition. Proc Natl Acad Sci U S A 102(30), 10493–10498. doi: 0501732102 [pii] 10.1073/pnas.0501732102.

Nicastro, G., Todi, S.V., Karaca, E., Bonvin, A.M., Paulson, H.L., and Pastore, A. (2010). Understanding the role of the Josephin domain in the PolyUb binding and cleavage properties of ataxin-3. PLoS One 5(8), e12430. doi: 10.1371/journal.pone.0012430.

Ohtake, F., Tsuchiya, H., Saeki, Y., and Tanaka, K. (2018). K63 ubiquitylation triggers proteasomal degradation by seeding branched ubiquitin chains. Proc Natl Acad Sci U S A 115(7), E1401–E1408. doi: 10.1073/pnas.1716673115.

Otten, E.G., Werner, E., Crespillo-Casado, A., Boyle, K.B., Dharamdasani, V., Pathe, C., et al. (2021). Ubiquitylation of lipopolysaccharide by RNF213 during bacterial infection. Nature 594(7861), 111–116. doi: 10.1038/s41586-021-03566-4.

Pao, K.C., Wood, N.T., Knebel, A., Rafie, K., Stanley, M., Mabbitt, P.D., et al. (2018). Activity-based E3 ligase profiling uncovers an E3 ligase with esterification activity. Nature 556(7701), 381–385. doi: 10.1038/s41586-018-0026-1.

Paulson, H.L. (2007). Dominantly inherited ataxias: lessons learned from Machado-Joseph disease/spinocerebellar ataxia type 3. Semin Neurol 27(2), 133–142.

Paulson, H.L., Perez, M.K., Trottier, Y., Trojanowski, J.Q., Subramony, S.H., Das, S.S., et al. (1997). Intranuclear inclusions of expanded polyglutamine protein in spinocerebellar ataxia type 3. Neuron 19(2), 333–344. doi: S0896-6273(00)80943-5 [pii],

Reina, C.P., Nabet, B.Y., Young, P.D., and Pittman, R.N. (2012). Basal and stress-induced Hsp70 are modulated by ataxin-3. Cell Stress Chaperones 17(6), 729–742. doi: 10.1007/s12192-012-0346-2.

Reina, C.P., Zhong, X., and Pittman, R.N. (2010). Proteotoxic stress increases nuclear localization of ataxin-3. Hum Mol Genet 19(2), 235–249. doi: ddp482 [pii] 10.1093/hmg/ddp482.

Scaglione, K.M., Zavodszky, E., Todi, S.V., Patury, S., Xu, P., Rodriguez-Lebron, E., et al. (2011). Ube2w and ataxin-3 coordinately regulate the ubiquitin ligase CHIP. Mol Cell 43(4), 599–612. doi: 10.1016/j.molcel.2011.05.036.

Schmidt, M., and Finley, D. (2014). Regulation of proteasome activity in health and disease. Biochim Biophys Acta 1843(1), 13–25. doi: 10.1016/j.bbamcr.2013.08.012.

Schmidt, T., Landwehrmeyer, G.B., Schmitt, I., Trottier, Y., Auburger, G., Laccone, F., et al. (1998). An isoform of ataxin-3 accumulates in the nucleus of neuronal cells in affected brain regions of SCA3 patients. Brain Pathol 8(4), 669–679.

Schols, L., Bauer, P., Schmidt, T., Schulte, T., and Riess, O. (2004). Autosomal dominant cerebellar ataxias: clinical features, genetics, and pathogenesis. Lancet Neurol 3(5), 291–304.

Seidel, K., Meister, M., Dugbartey, G.J., Zijlstra, M.P., Vinet, J., Brunt, E.R., et al. (2012a). Cellular protein quality control and the evolution of aggregates in spinocerebellar ataxia type 3 (SCA3). Neuropathol Appl Neurobiol 38(6), 548–558. doi: 10.1111/j.1365-2990.2011.01220.x.

Seidel, K., Siswanto, S., Brunt, E.R., den Dunnen, W., Korf, H.W., and Rub, U. (2012b). Brain pathology of spinocerebellar ataxias. Acta Neuropathol 124(1), 1–21. doi: 10.1007/s00401-012-1000-x.

Sittler, A., Muriel, M.P., Marinello, M., Brice, A., den Dunnen, W., and Alves, S. (2017). Deregulation of autophagy in postmortem brains of Machado-Joseph disease patients. Neuropathology, doi: 10.1111/neup.12433.

Tan, J.M., Wong, E.S., Kirkpatrick, D.S., Pletnikova, O., Ko, H.S., Tay, S.P., et al. (2008). Lysine 63-linked ubiquitination promotes the formation and autophagic clearance of protein inclusions associated with neurodegenerative diseases. Hum Mol Genet 17(3), 431–439. doi: 10.1093/hmg/ddm320.

Toulis, V., Garcia-Monclus, S., de la Pena-Ramirez, C., Arenas-Galnares, R., Abril, J.F., Todi, S.V., et al. (2020). The Deubiquitinating Enzyme Ataxin-3 Regulates Ciliogenesis and Phagocytosis in the Retina. Cell Rep 33(6), 108360. doi: 10.1016/j.celrep.2020.108360.

Winborn, B.J., Travis, S.M., Todi, S.V., Scaglione, K.M., Xu, P., Williams, A.J., et al. (2008). The deubiquitinating enzyme ataxin-3, a polyglutamine disease protein, edits Lys63 linkages in mixed linkage ubiquitin chains. J Biol Chem 283(39), 26436–26443.

Yang, C.S., Jividen, K., Spencer, A., Dworak, N., Ni, L., Oostdyk, L.T., et al. (2017). Ubiquitin Modification by the E3 Ligase/ADP-Ribosyltransferase Dtx3L/Parp9. Mol Cell 66(4), 503–516 e505. doi: 10.1016/j.molcel.2017.04.028.

